# Tagging and harvesting of developing rice spikelets to demarcate grain developmental stages

**DOI:** 10.1101/2025.10.15.682554

**Authors:** Antima Yadav, P V Aswathi, Upasana Das, Falah Qasim, Tripti Avinash, Pinky Agarwal

## Abstract

**Premise:** In rice, grain is the major sink for nutrient storage and is a rich source of carbohydrates, seed storage proteins, lipids, minerals, and vitamins, making it a principal food source for over half of the world’s population. Grain development in rice generally spans over a month and has been divided into five stages, S1-S5, depending upon various morphological and physiological changes occurring in the developing seed. To completely understand the process, rice grain development needs to be examined at each stage. However, there are no detailed protocols available to demarcate grain developmental stages.

**Methods and Results:** This protocol describes how to tag and harvest the developing rice spikelets. The process starts with anthesis and continues till seed maturation, demarcating the five stages of rice grain development. Seeds are collected, labelled, and stored for use in molecular experiments. Using this protocol, we were able to identify and distinguish different seed developmental stages as observed by histochemical screening.

**Conclusion:** This is an easy, efficient, and cost-effective method. This protocol ensures the collection of the entire panel of rice seed developmental stages. The stored tissue can later be used for various molecular biology based experiments.

## INTRODUCTION

Rice seed not only acts as precursor for the new generation but also is a major source of nutrition for over half of the world’s population. Global demand for rice is continuously increasing because of the rising population. As the expansion of rice cultivated land is unlikely, an acceleration in overall rice yield is necessary to feed the growing population (Arif et al., 2019). Flowering becomes important to study because it affects grain filling and subsequently, yield in rice (Lu et al., 2022; Mishra et al., 2022). There have been ample studies which have focused on rice flowering from various directions, like, anthesis and fertilization (Shi et al., 2018), crop improvement by breeding and genetics (Hori et al., 2016; Wang et al., 2020b; Lin et al., 2021; Pruthi et al., 2022) and response to different stress conditions (Cho et al., 2017; Wu et al., 2022). The vegetative and reproductive phase of rice development is very sensitive and highly influenced by environmental conditions. Multiple studies have shown the effect of different stresses on rice flowering and its overall development (Wang et al., 2020a; Mishra et al., 2022). Therefore, division and demarcation of developmental stages, both vegetative and reproductive, is of utmost importance. This demarcation enables us to identify an appropriate developmental stage and hence, we can compare phenotypes of different varieties or stress conditions. Rice seed development spans over a month, starting from anthesis till a mature rice grain is formed. Rice grain formation starts at the onset of a double fertilization event in which one male gamete fuses with the egg cell (syngamy) giving rise to an embryo and the other male gamete fertilizes the secondary nucleus (triple fusion) resulting in the formation of the endosperm (Guignard, 2001; Kordium, 2008, Agarwal et al., 2011). Seed development in rice has been divided into five stages (S1-S5) based on morphological and physiological changes occurring in the embryo and the endosperm (Agarwal et al., 2011). To collect seeds of different seed developmental stages, panicles are tagged on the day of anthesis and this day is referred to as 0 days after pollination (DAP). Panicle emergence should be closely examined, as the entire protocol starts with panicle emergence. Spikelets tagged between 0-2 DAP fall in S1 stage. During this stage, rapid cell division takes place in the embryo resulting in a globular embryo, however, the endoperm forms a syncytium. 3-4 DAP represents S2 stage of seed development. During this stage, organ initiation happens in the embryo and endosperm cellularizes, resulting in a green coloured, liquid state endosperm. The next stage is S3, which spans from 5-10 DAP. During this stage organ enlargement proceeds, forming completely developed organs. Endosperm is starchy and in milky stage. S4 stage covers 11-20 DAP. A completely developed embryo possesses SAM, coleoptile, scutellum, radicle, and epiblast. By 20 DAP embryo is mature and enters the dormant stage. For endosperm, this stage serves as the storage phase involving starch synthesis and storage protein accumulation. Endosperm transits from soft dough stage to hard dough stage. During the final stage (S5; 21-29 DAP), maturation of the seed takes place, giving rise to mature rice grain which is majorly occupied by the endosperm and enclosed by the husk (Agarwal et al., 2011). Seeds at different developmental stages are undergoing different processes which are regulated by various molecular networks (Cai et al., 2019; Zhou et al., 2021). To understand these molecular events which contribute to rice seed development, it becomes essential to demarcate the different seed developmental stages. Our protocol aims at delineating rice seed development in such a way that all the changes occurring during the process are thoroughly covered. In our protocol, we will demonstrate how the rice spikelets are identified for tagging and after identification, panicles are tagged. For harvesting, seed stage is inferred according to DAP. Post-harvest frozen tissues are stored at -80° C.

## METHODS AND RESULTS

### Tagging to demarcate the seed developmental stages

Varying environment largely influences seed development processes. To avoid wastage of time and resources and to ensure accurate data collection, researchers need to know about how rice seed development occurs. The critical timing of data collection is often determined too late, resulting in constricted time interval for sampling. We have developed an easy way to tag the rice spikelets, which allows precise data collection and efficient use of tissue samples during the off-season. This is an organized method to categorize the five stages of rice seed development. This technique helps us to avoid mistakes during sample collection and ensures collection of an entire panel of seed development stages. This method of sampling is utilized to keep the tissues intact for various analyses such as enzyme assays, quantification, gene expression, starch, protein, and lipid analysis.

The development and initiation of the panicle is the starting point of the reproductive growth phase. The primary and secondary branches of the panicle differentiate within the flag leaf sheath. It is not visible until it exerts from the boot. At this time the tip of the panicle is above the collar of the flag leaf. The period between this day and the following day is a critical time for tagging because, during this time, the anthers and filaments emerge from the florets on the panicle. Anthesis occurs just after the florets open up and are exposed to light and air. This exposure allows the anther and filament to protrude out of the floret, which enables pollination and fertilization to occur (Yoshida and Nagato, 2011).

Before starting the actual tagging protocol, it is very essential to examine the developmental stage of the rice plant. Before panicle emergence, by hand touching and feeling the thickened flagleaf sheath, the embedded panicle development has been confirmed. After the leaf sheath thickening, we visited the plants everyday as it takes 1-2 days for the emerged panicle to anthesis (Figure 1A and B). In rice, anthesis starts from 6 am till 11 am. As and when the panicle emerges, the first step was to make tags. To make a tag, a colourful label of approximately 3-4 cm were cut and mentioned the date of anthesis using a water-proof marker (Figure 2). Lastly, covered the label using a cello tape. Pollination in rice happens in a basipetal manner. During pollination/anthesis, spikelet opens, followed by elongation of the anther filament. This can be easily visualised with off white anthers coming out of an open spikelet (fresh anthers) (Figure 1B). The panicles were tagged by putting the tag at the base of 2-3 panicle branches together, above which dehisced anthers were present. Utmost care was taken while tagging to choose healthy panicles and not to disturb the anthers as it may affect the fertilization. Tagging on the same panicle continued depending upon the emergence of fresh anthers. For example, a panicle was tagged on 13.10.2024 at the base till where the dehisced anthers were visible. Next day some fresh anthers were seen below the 13.10.2024 tag, then a 14.10.2024 tag was put on the lower portion up to the region where fresh anthers were visible. Flowering in a rice field is completed in around 14 days, 8-10 panicles were tagged each day until the fresh anthers stopped emerging from the primary tillers. One single panicle was not tagged for more than 3 days continuously, as the inferior spikelets often develop slowly or do not pollinate at all.

**Figure 1.**
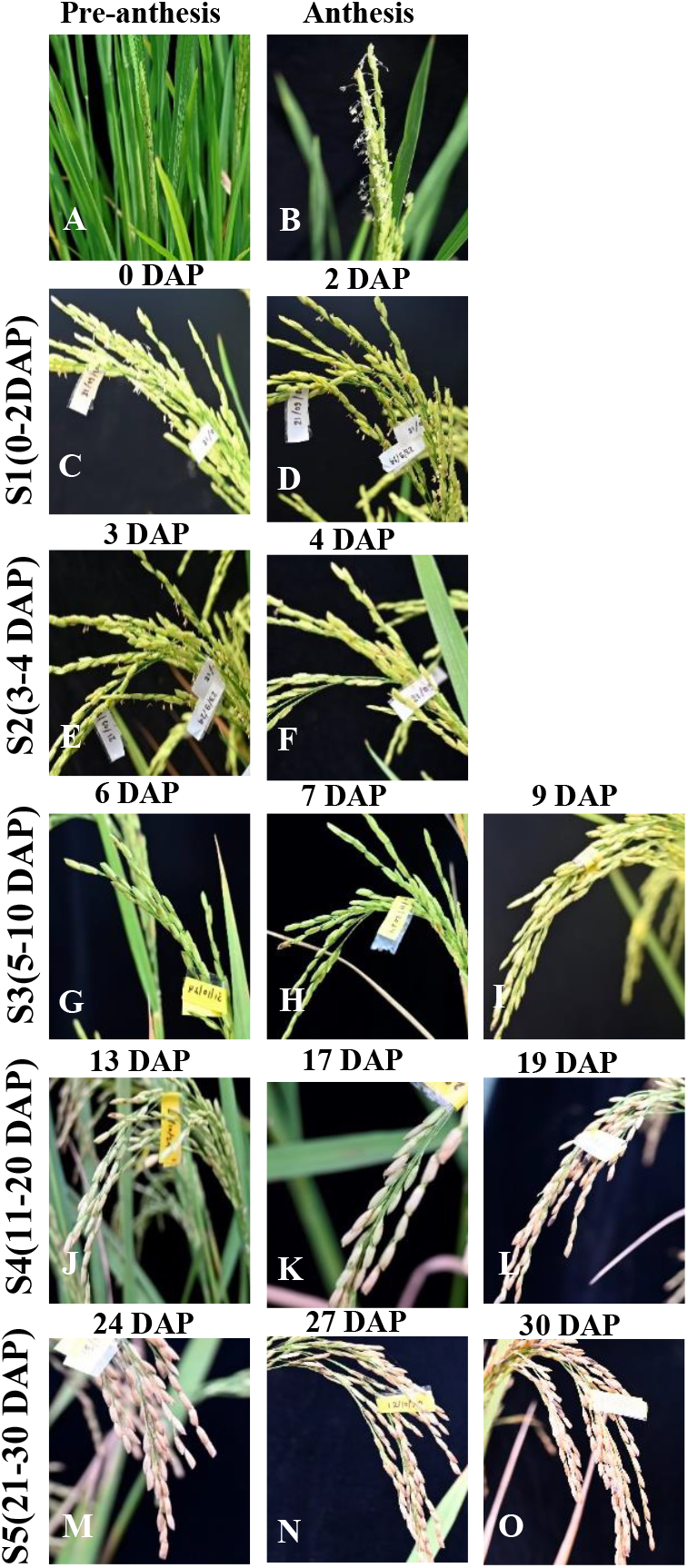
Stages of rice seed development from spikelet emergence to grain maturation. (A) Visual depiction of a pre-anthesis rice panicle. (B) Panicle at anthesis stage, characterized by the emergence of off-white anthers from spikelets. (C-D) Stage S1 of seed development (0-2 DAP), with spikelets labelled according to their date of flowering. (E-F) Stage S2 (3-4 DAP), representing early post-fertilization development. (G-H) Stage S3 (5–10 DAP), showing progressing grain filling. (I-J) Stage S4 (11–20 DAP), indicating active endosperm development and cellular differentiation. (M-O) Stage S5 (21–29 DAP), marking late grain filling and maturation.

**Figure 2.**
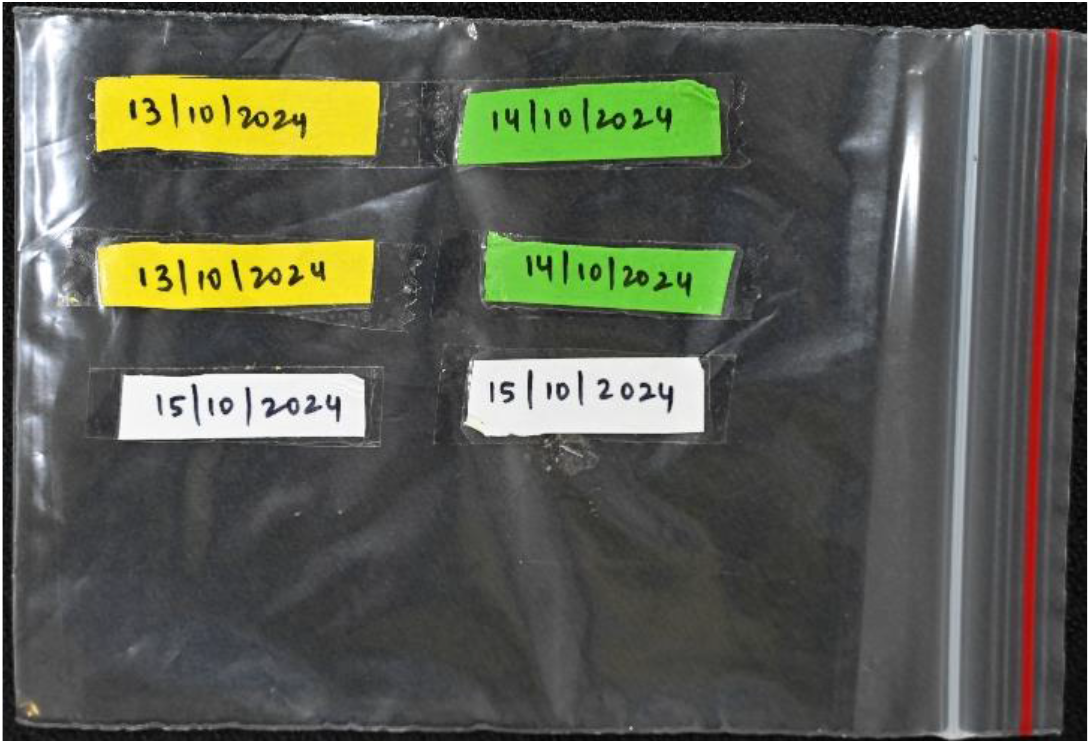
**Representation of a tag used to demarcate pollinated florets.**

### Harvesting and storage of sample

For harvesting, field bags were prepared before going to the field, which contained surface sterilized scissors, aluminium foils, marker pens, gloves and liquid nitrogen in a container. The samples were harvested using scissors, packed in aluminium foil pouches and collected in liquid nitrogen. The outer surface of aluminium foil was labelled in prior, keeping in mind the plant number/ replicate number and development stage of the seed sample that was to be collected. If carrying liquid nitrogen was not feasible, we collected the samples on ice to avoid degradation of various biomolecules.

Panicle tagging and harvesting goes hand in hand especially for the initial stages of seed development (S1, S2 and initial DAPs of S3) (Figure 1C-I). So, panicles were tagged in the morning and harvested according to DAP in the evening. Harvesting or sample collection were done in the morning or evening hours before watering the field. For calculating DAP, the date on which panicle was tagged is considered as 0 DAP and accordingly different DAPs were calculated. Harvesting dates were planned according to the stage at which seed samples have to be collected. For instance; if tagging was done on 12.10.2024, the seed samples for 3 DAP were collected on 15.10.2024. The chosen panicle was hold carefully without disturbing the rest of the panicles and cut the seed samples above the tag using scissors. Alternately, we also plucked enough number of spikelets from the tagged panicle to avoid the wastage of the entire panicle. While collecting S3-S5 (5-29 DAP) seed samples (Figure 1G-O), care was taken to avoid unfertilized/unfilled/shrivelled seeds. Seed filling was ensured by lightly pressing the immature seeds. Milky exudate from the seeds indicated normal grain filling. Later stages were identified by the colour change of husk from green to light brown and brown (Figure 1J-O).

In case of a multi-tagged panicle, if scissor cuts were made to collect a small portion of the panicle, the remaining panicle was not used for further collection, as the cut may cause stress and induce biochemical changes in the panicle. Generally, to collect samples from multi-tagged panicles, we planned the harvest in prior according to the dates on the tags. For example, a panicle that had tags of 13.10.2024 and 14.10.2024, the harvest was planned on 16.10.2024 to collect samples of 3 and 2 DAP, respectively. After the completion of sample collection, all the samples were distributed into cryoboxes labelled according to the development stages and stored at -80°C for long term usage.

### Downstream applications

Using this technique various scientific questions related to rice seed development from anthesis to seed maturation can be addressed in different rice varieties as well any mutants. Samples of different seed developmental stages can be used in the following experiments:

1. Cytological study – Samples of sequential development stages can be used to study seed development at cellular level (Table1). This has helped to understand the various steps in endosperm development; and the points which can be targeted for increment of grain size.
2. Expression analyses – to understand the expression pattern of various genes involved in the seed development (Table1). This has aided in understanding the genes and pathways controlling grain development, grain size, starch and seed storage protein biosynthesis, which can be further modified using genetic engineering techniques.
3. Starch estimation and analyses – to study the starch synthesis and accumulation in relation to seed development (Table1). This helps to understand the pattern of starch production, and in relation with expression analyses, has been used to knock out or over express genes to increase amylose or amylopectin content, as desired.
4. Protein estimation and analyses – to investigate the seed development process at protein level as well as to get detailed understanding about storage proteins (Table1). Rice is a major protein source in many South-East Asian countries, as it is an inexpensive staple food. Understanding the production of seed storage proteins can aid in increasing the total grain protein content, and hence target malnutrition.
5. Metabolome study – to unravel the relation between genetic and metabolic pathways in determining seed related traits (Table1). Limited studies are available which decipher the role of various metabolites in grain development. Metabolome studies can fill this gap.
6. Transcriptome/ Epitranscriptome study – transcriptome study is a powerful tool to investigate and identify a subset of genes active at specific point of time in the seed tissue, to get a clearer picture from seed formation to maturation (Table 1). Such genes can then be tweaked to generate rice plants with a desirable phenotype, targeting crop improvement. Epitranscriptome studies on seed developmental stages allow to identify dynamic chemical modifications happening in the RNA transcripts (coding and non-coding).
7. Sequencing – stage specific samples can be used for various sequencing techniques such as genetic sequencing, chromatin immuno-precipitation (ChIP) sequencing etc. to delineate the molecular basis of seed development. This will unravel various networks operable during the process of grain formation, which can then be targeted as per the need.
8. Epigenetic studies can be performed to understand the influence of histone modifications like methylation, acetylation, and deacetylation etc. on gene expression in different seed developmental stages, to deepen our knowledge on the complex process of rice grain development.

**Table 1.**
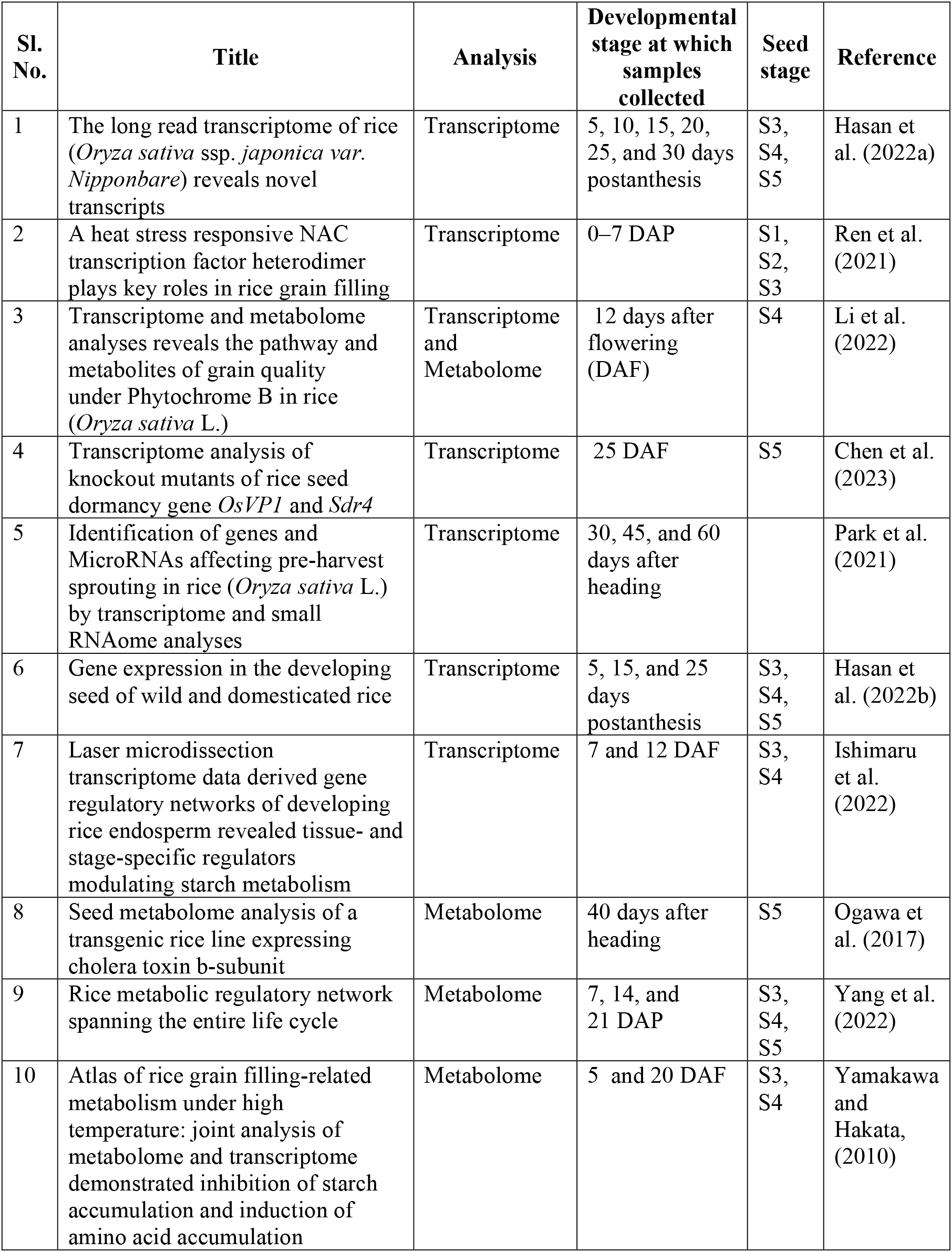
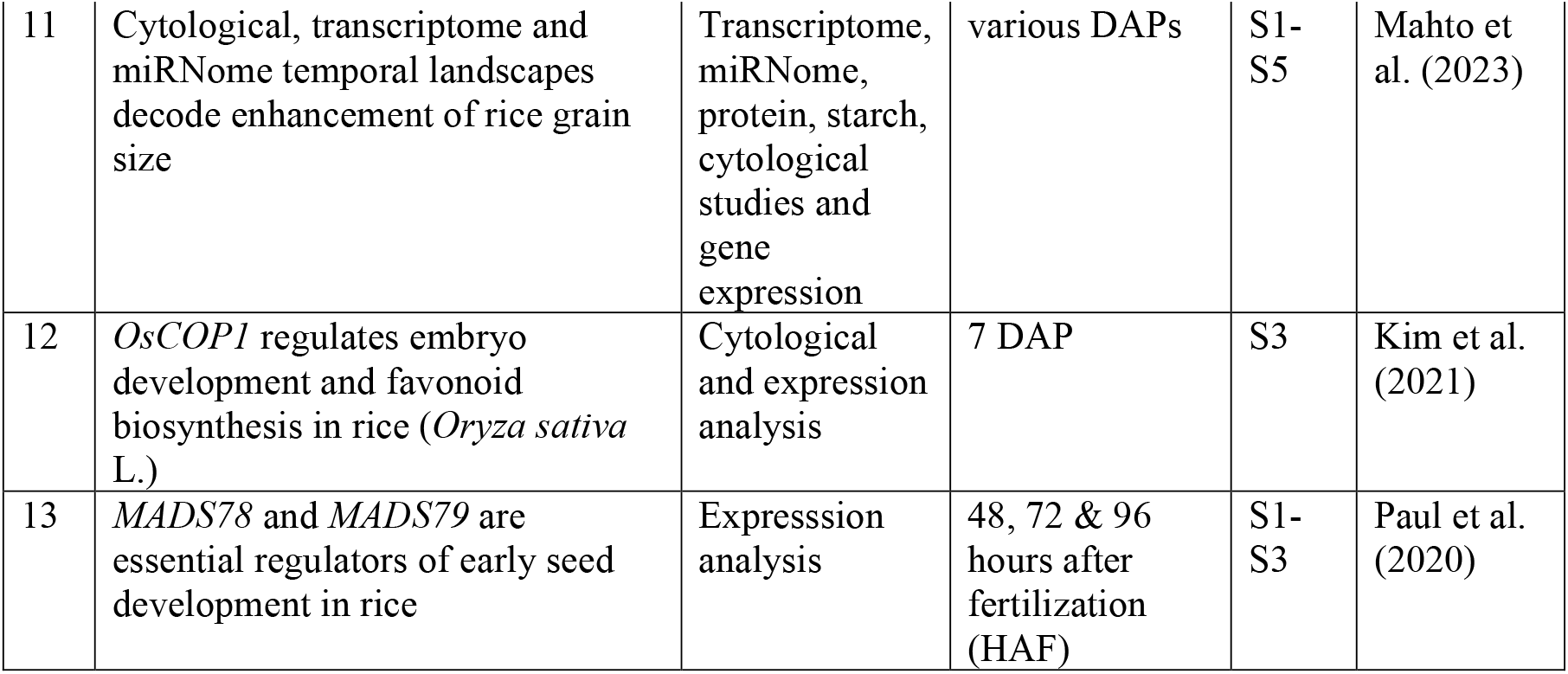
Studies with downstream applications.

## CONCLUSIONS

This method for tagging developing rice spikelets is more organized and user-friendly than traditional sampling methods. In a protocol described for tracking the seed development stages (Counce and Moldenhauer, 2019), the culm of the panicle is identified by observing the emergence of the final leaf or flag leaf, and marked with a plastic numbering tag and waterproof ink. Due to the uneven dehiscence of anthers and the pollination followed by it, the exact tracking of stages will not be precise. Here, the tagging of the rice spikelets by examining the changes in the panicle would be more comprehensible. There are automated methods such as detecting flowering panicles of rice using field-acquired time-series RGB images (Guo et al., 2015). These are more expensive and there will be a need of technical assistance for the equipments used. Compared to these, we have arrived at this protocol through a process of trial and error, after several years of observation and study on rice reproductive development.

## ACKNOWLEDGMENTS

The authors are thankful to the University Grants Commission for Ph. D. research fellowship; Department of Biotechnology, and Department of Science and Technology, Ministry of Science and Technology, India for grants supporting the research in this paper; and to NIPGR for core grant. The authors acknowledge the support provided by DBT e-library consortium for providing online access to articles. PA acknowledges the grants SPG_2021_002899 and CRG_2018_000501 from the Science and Engineering Research Board (SERB), Ministry of Science and Technology, India.

## AUTHOR CONTRIBUTIONS

PA conceived and A. Y., A. P. V., F. Q., U. D., T. A., A. M. conducted the study. P. A., A. Y., A. P. V. and S. K. drafted the manuscript. All authors contributed to and approved the final manuscript.

## DATA AVAILABILITY STATEMENT

No supporting data were used.

